# Cardiac gene therapy with PDE2A limits remodeling and arrhythmias in mouse models of heart failure induced by catecholamines

**DOI:** 10.1101/2023.04.17.537274

**Authors:** Rima Kamel, Aurélia Bourcier, Jean Piero Margaria, Audrey Varin, Agnès Hivonnait, Françoise Mercier-Nomé, Delphine Mika, Alessandra Ghigo, Flavien Charpentier, Vincent Algalarrondo, Emilio Hirsch, Rodolphe Fischmeister, Grégoire Vandecasteele, Jérôme Leroy

## Abstract

**BACKGROUND:** Constitutive cardiac PDE2 activation was shown to protect against contractile dysfunction and arrhythmia in heart failure (HF). However, it remains unknown whether an acute elevation of PDE2 is efficient to prevent maladaptive remodeling and arrhythmia. In this study we tested whether increasing acutely PDE2A activity in preclinical models of HF using cardiac PDE2 gene transfer could be of therapeutic value.

**METHODS AND RESULTS:** C57BL/6 male mice were injected with serotype 9 adeno-associated viruses (AAV9) encoding for PDE2A, or luciferase (LUC). Cardiac function assessed by echocardiography unveiled neither structural change nor dysfunction consecutive to PDE2A overexpression while AAV9 inoculation led to a ≈10-fold rise of PDE2A protein levels. Two weeks after AAV9 injections, mice were implanted with osmotic minipumps delivering NaCl or isoproterenol (Iso) (60 mg/kg/day) or Iso and phenylephrine (Iso+Phe, 30 mg/kg/day each) for 2 weeks. In LUC mice, chronic infusion with Iso increased left ventricular (LV) weight over body weight ratio, promoted fibrosis and decreased ejection fraction, but animals overexpressing PDE2A were protected towards these deleterious effects. Similarly, concomitant treatment with Iso+Phe promoted LV contractile dysfunction, fibrosis and apoptosis in LUC mice, while PDE2A overexpression limited these adverse outcomes. Furthermore, inotropic responses to Iso of ventricular cardiomyocytes isolated from Iso+Phe-LUC mice loaded with 1 µmol/L Fura-2AM and stimulated at 1 Hz to record calcium transients and sarcomere shortening were dampened. Chronic treatment with catecholamines favoured spontaneous calcium waves upon β-AR stimulation at the cellular level and promoted susceptibility to ventricular arrhythmias *in vivo* evoked by catheter-mediated ventricular pacing after Iso and atropine injection. However, these adverse effects were blunted by the cardiac gene therapy with PDE2A.

**CONCLUSION:** Gene therapy with PDE2A limits cardiac adverse left ventricle remodeling and dysfunction induced by catecholamines as well as ventricular arrhythmias, providing evidence that acutely increasing PDE2A activity could prevent progression towards HF.

## INTRODUCTION

The “fight-or-flight” response is launched by a discharge of norepinephrine by the sympathetic nervous system, amplified by the secretion of catecholamines by the adrenal medulla which act on cardiomyocytes via the β-adrenergic receptors (β-AR)/adenylyl cyclase (AC)/cAMP/protein kinase A (PKA) axis to stimulate cardiac rhythm (chronotropy), contractile force (inotropy) and relaxation (lusitropy).^1^ PKA phosphorylates many of the proteins critically involved in the excitation-contraction coupling (ECC). These include the small G protein Rad to stimulate the L-type Ca^2+^ channels (LTCCs),^2^ the ryanodine receptor 2 (RyR2) to increase its open probability, the phospholamban (PLB) to relieve its inhibitory effect on the sarco/endoplasmic reticulum Ca^2+^-ATPase (SERCA2) and contractile proteins, such as troponin I (TnI) and myosin binding protein C.^1^ While this acute β-AR activation is favorable allowing the adaptation of the cardiac output to increased body’s metabolic demand during a stress or an effort, a chronic overactivation of the sympathetic nervous system is deleterious. In heart failure (HF), the decreased cardiac function leads to a chronic catecholamine elevation^3^ which elicits maladaptive remodeling comprising cardiac hypertrophy, cardiomyocyte death, cardiac fibrosis and promotes arrhythmias thus aggravates pump dysfunction. The excessive β-AR stimulation is central in this propelled vicious circle as attested by the beneficial effects of β-adrenergic antagonists which are the first-line therapy to treat chronic HF with reduced ejection fraction.^4, 5^ However, despite an enhancement in survival rates by the use of β-blockers,^5–7^ the mortality of HF patients 5 years after diagnosis mainly due to pump failure or ventricular arrhythmias remains unacceptably high at ∼50%.^8^ Furthermore, in many patients, intolerance does not allow titration to an effective dosage.^9^ There is thus a need for new therapeutic approaches to treat HF.

Intracellular cAMP levels produced upon β-AR stimulation are counterbalanced by the degradation of the cyclic nucleotide by enzymes called phosphodiesterases (PDEs) which not only terminate the activation of cAMP effectors but also compartmentalize this second messenger in discrete subcellular compartments.^10^ Among the 6 PDE families expressed in cardiomyocytes to degrade cAMP, PDE2 has been recently identified as a therapeutic target to treat HF. PDE2 was shown to inhibit cardiac LTCC activity in various species, including humans^11^ and to control ECC.^12–14^ In contrast to PDE3 and PDE4 which expression and activities are generally decreased in pathological hypertrophy and HF,^15–17^ PDE2 is increased in HF to blunt β-AR/cAMP signals suggesting a protective mechanism against increased circulating catecholamines.^18^ Accordingly, transgenic (TG) mice with constitutive cardiac overexpression of PDE2 exhibit improved cardiac function and are protected against catecholamine-induced arrhythmias after ischemic injury.^19^ Moreover, PDE2 hydrolytic activity is stimulated by 10 to 30-fold by cGMP via binding to the regulatory N-terminal GAF-B domain^20^ making PDE2 a unique integrator of cAMP/cGMP signals^12, 21, 22^. Interestingly, a recent study unveiled that the antiarrhythmic effects of CNP occur via PDE2 stimulation, attenuating spontaneous Ca^2+^ events, limiting CaMKII activation and decreasing the late Na^+^ current promoted by a β-AR stimulation.^23^ However, cardioprotective effects of increased PDE2 activity is a matter of debate since conflicting results reported that PDE2 upregulation is pro-hypertrophic.^24^ Furthermore, its inhibition was found to antagonize cardiomyocyte growth,^24^ to protect against apoptotic cell death by promoting mitochondrial elongation^25^ and PDE2 blockade with BAY60-7550 hindered cardiac maladaptive remodeling by enhancing NO/GC/cGMP signaling.^26^

In an attempt to further delineate the impact of PDE2 upregulation on cardiac function and its putative beneficial effects in HF, we performed gene therapy with adeno-associated viruses serotype 9 (AAV9) allowing preferential cardiac overexpression of PDE2A3. This strategy allows expression from a definite time point rather than constitutive cardiac expression as previously employed.^19^ In healthy mice transduced with AAV9-PDE2A3, the 10-fold increase in PDE2 expression obtained did not affect cardiac function and did not evoke hypertrophy. On the contrary, it lowered left ventricle (LV) hypertrophy elicited by a chronic exposure to catecholamines, counteracted their pro-fibrotic effects and limited LV dysfunction. At the cellular level, gene therapy with PDE2A3 improved ECC responsiveness to β-AR, increasing Ca^2+^ transient and contraction amplitude upon isoproterenol, suggesting less β-AR desensitization. Furthermore, while polymorphic ventricular tachycardias were triggered using catheter-mediated ventricular pacing in LUC mice treated chronically with catecholamines, none of the animals injected beforehand with AAV9-PDE2A3 exhibited arrhythmias. Thus, myocardial increase of PDE2A activity limits cardiac maladaptive remodeling and dysfunction induced by catecholamines as well as associated ventricular arrhythmias, further demonstrating that it could constitute an alternate or a complementary anti-adrenergic therapeutic approach to β-blockers to treat HF.

## METHODS

### Ethical approval and animals

All procedures were carried out at the animal facility and at the laboratory INSERM UMR-S1180 of the faculty of Pharmacy of the Paris-Saclay University, according to institutional regulations and the local guide for the use and care of laboratory animals. All experiments involving animals were conducted in agreement with the European Community guiding principles in the welfare and use of animals (2010/63/UE) and the French decree no. 2013-118 on the protection of animals used for scientific purposes. The protocol was approved by the local ethics committee (CREEA Ile-de-France Sud) and by the French Ministry of higher Education and Research (17852-2015051112125554 v4). In this study, 8-week-old male C57BL/6 mice were used.

### Reagents

Isoproterenol (Iso), phenylephrine (Phe) and atropine were purchased from Sigma-Aldrich (St. Louis, Missouri, USA). Bay60-7550 was purchased from Cayman chemical (Ann Arbor, Michigan, USA).

### AAV9-LUC/PDE vectors production, purification, characterization and injection

AAV9-PDE2A holds a PDE2A expression cassette flanked by two AAV2 inverted terminal repeats. The expression sequence is pseudotyped with an AAV9 capsid. The cassette contains a CMV promoter, a β-globin intron, a FLAG tag fused in 5’ of the complete murine PDE2A3 coding sequence (NCBI: NM_001008548.4) and an hGH polyadenylation signal. AAV9-PDE2A was produced in AAV-293T cells (Stratagene #240073, San Diego, California, USA) with the three-plasmid method and the calcium-chloride transfection. The virus was purified by cesium-chloride gradient. A RT-PCR on the human cytomegalovirus (CMV) promoter was used to determine viral particles titers. An expression cassette encoding firefly luciferase (LUC) was used for control condition. A CMV promoter controlling luciferase expression, was packaged into AAV9 capsids and purified on cesium-chloride gradient to yield AAV9-LUC virus. AAV9-LUC or AAV9-PDE2A were injected *via* the tail vein at 10^12^ viral particles/mouse. AAV9-CMV is the empty control vector used to produce AAV9-PDE2A containing solely the CMV promoter without any open reading frame sequence. The AAV9-PDE4B has been previously described.^17^

### Transthoracic echocardiography

Transthoracic two-dimensional-guided M-mode echocardiography of mice was performed using an echocardiograph with a ML6 linear probe of 15 MHz (Vivid E9, General Electric Healthcare®, Chicago, Illinois, USA) under 1-2% isoflurane gas and 0.4-0.8 L/min oxygen anesthesia. For isoflurane induction, mice were placed for approximately 2 min in an isolation chamber filled with isoflurane (5% in 100% oxygen) at a flow rate of 0.5–1 L/min. Body temperature was maintained at 37°C by the use of a thermally controlled heating pad (Harvard Apparatus). Measurements from animals with heart frequency bellow 400 bpm were discarded. LV cavity and wall thickness dimensions during systole and diastole were determined using two parasternal axes: short and long. The left ventricular mass (LVM) was calculated according to the Penn formula while assuming a spherical LV geometry and validated for the mouse heart (LVM=1.04×[(LVIDd+IVS+PW)^3^−(LVIDd)^3^], where 1.04 is the specific gravity of muscle. LVIDd: Left Ventricular Internal Diameter in diastole, IVSd and PWd are end-diastolic InterVentricular Septum and Posterior Wall thicknesses, respectively. Analysis was performed for the echocardiographic images using the EchoPac software (General Electric Healthcare®). For each of the two axes and the measured parameters, three measurements on three different sections were averaged.

### Catecholamines infusion models

Two weeks after AAV9 injections, mice were treated for 14 days with either isoproterenol (60 mg/kg/day), isoproterenol+phenylephrine (30 mg/kg/day each) or vehicle (0.9% NaCl) which were administered via osmotic minipumps (2002, Alzet®, Cupertino, California, USA) surgically implanted subcutaneously in the back of the mice as previously described.^17^ After two weeks, minipumps were removed. Animals were either sacrificed following minipumps removal or two weeks later for the isoproterenol model.

### Morphological and histological analysis

For euthanasia, mice were anesthetized by an intraperitoneal injection of Doléthal® (100 mg/kg). Afterwards, hearts were rapidly excised and remaining blood was washed out in cold Ca^2+^-free phosphate buffer saline solution (155.2 mmol/L NaCl, 1.06 mmol/L KH_2_PO_4_ and 2.9 mmol/L NaH_2_PO_4_). Hearts and Lungs were weighed and tibia length was measured. For histology, a transversal slice of 3-4 mm width was cut in the middle of the heart and rapidly fixed in 4% paraformaldehyde. The rest of ventricular tissue was frozen in liquid nitrogen and stored at −80°C until use. Histological staining was performed using a standard protocol. After fixation of the hearts for 24 h in 4% paraformaldehyde, they were paraffin embedded and transversely sectioned. Sections (5 or 7 μm) were deparaffinized. Afterwards, samples were subjected to Heat-Induced Epitope Retrieval (HIER) in a 10 mmol/L citrate buffer, pH=6. A Masson’s trichrome stain kit (Microm, Brignais, France) was used to assess cardiac fibrosis. To quantify apoptotic myocytes in the heart, autofluorescence was quenched by treating paraffin-embedded sections with PBS/BSA (5%) for 2 h before performing TUNEL staining (Roche, Basel, Switzerland) according to the manufacturer’s protocol and using Proteinase K treatment. Sections were counterstained with Alexa Fluor 594-conjugated wheat germ agglutinin (WGA, Invitrogen, Carlsbad, California, USA) at 10 μg/mL for 1 h at room temperature to visualize cell membranes. Slides were scanned by the digital slide scanner NanoZoomer 2.0-RS (Hamamatsu, Japan), which allowed an overall view of the samples. Images were digitally captured from the scan slides using the NDP.view2 software (Hamamatsu, Japan).

### ECG recording and intracardiac recording and pacing

The criteria used to measure RR, PR, QRS and QT intervals on recorded ECG have been described elsewhere.^27^ The QT interval was corrected for heart rate with the Bazett formula adapted to mouse sinus rate, *i.e.* QTc = QT/(RR/100)1/2 with QT and RR, expressed in ms.^28^ Mice were anesthetized with an injection of etomidate of 28 mg/kg (Hypnomidate^®^ 2 mg/ml, Janssen-Cilag, Issy les Moulineaux, France) before and after intracardiac and surface ECG recording and pacing. The extremity of a 2F octapolar catheter (Biosense Webster) was placed in the right ventricle through the right internal jugular vein, using intracardiac electrograms as a guide for catheter positioning in the right ventricle. Surface ECG (lead I) and intracardiac electrograms were recorded on a computer through an analog-digital converter (IOX 1.585, Emka Technologies, Paris, France) for monitoring and later analysis (ECG-Auto v3.3.0.5, Emka Technologies, Paris, France). Intracardiac electrograms were filtered between 0.5 and 500 Hz. Pacing was performed with a digital stimulator (DS8000, World Precision Instruments, Sarasota, Florida, USA). Standard pacing protocols were used to determine the ventricular effective refractory periods (VERP) and to induce ventricular arrhythmias. Determination of stimulation threshold, which corresponds to the minimal energy provoking a ventricular depolarization, is realized with an impulse duration at 1 ms and with an increasing intensity. Afterwards, stimulations were conducted at intensities 1.5 times the excitation threshold. VERP were assessed at baseline by using the programmed electrical stimulation (PES) method. Extrastimuli were delivered following trains of 20 impulses at a cycle length of 70 ms followed by one, two and then three extrastimuli. The extrastimulus coupling interval was initially set at 70 ms and then reduced by 2 ms at each cycle until ventricular refractoriness was reached. The inducibility of ventricular arrhythmias was assessed in baseline conditions and 3 min after intraperitoneal injection of Iso 1.5 mg/kg and atropine 1 mg/kg. Ventricular tachycardias were defined as the occurrence of at least 4 consecutive QRS complexes with a morphology different than that seen with a normal sinus rhythm, after last stimulated beat.

### Isolation of mouse cardiomyocytes

Mice were anesthetized by intraperitoneal injection of Doléthal^®^ (100 mg/kg). The heart was quickly removed and placed into a cold Ca^2+^-free Tyrode’s solution containing 113 mmol/L NaCl, 4.7 mmol/L KCl, 1.2 mmol/L MgSO_4_-7H_2_O, 0.6 mmol/L KH_2_PO_4_, 0.6 mmol/L NaH_2_PO_4_, 1.6 mmol/L NaHCO_3_, 10 mmol/L HEPES, 30 mmol/L taurine, 20 mmol/L glucose and 17.03 µmol/L insulin adjusted to pH 7.4. The ascending aorta was cannulated, and the heart was perfused with oxygenated Ca^2+^-free Tyrode’s solution at 37°C for approximately 4 min using retrograde Langendorff perfusion. Subsequently, the heart was perfused with Ca^2+^-free Tyrode’s solution containing Liberase^TM^ research grade (Roche Diagnostics, Basel, Switzerland) for 10 min at 37°C for enzymatic dissociation. Afterwards, the heart was removed and placed into a dish containing a Tyrode’s solution supplemented with 0.2 mmol/L CaCl_2_ and 5 mg/ml BSA (Sigma-Aldrich, St. Louis, Missouri, USA). Atria were discarded and ventricles were cut into small pieces, and triturated gently with a pipette to disperse the myocytes. Ventricular myocytes were filtered on gauze and allowed to sediment by gravity for approximately 7 min. The supernatant was removed, and cells were suspended in Tyrode’s solution supplemented with 0.5 mmol/L CaCl_2_ and 5 mg/ml BSA. Sedimentation was repeated and cells were finally suspended in a Tyrode’s solution with 1 mmol/L CaCl_2_. For Ionoptix experiments, freshly isolated ventricular myocytes were plated for 45 min at a density of 10^4^ cells per dish in 35-mm culture dishes coated with laminin (10 μg/ml) and stored in a cell incubator at 37°C, 5% CO_2_ until use.

### Calcium transients and sarcomere shortening measurements

Isolated cardiomyocytes were loaded with 1 μM Fura-2 AM (Invitrogen, Carlsbad, California, USA) at room temperature for 15 min and then washed for 15 additional min with an external Ringer solution containing (in mmol/L): NaCl 121.6, KCl 5.4, NaHCO_3_ 4.013, NaH_2_PO_4_ 0.8, 10 mM HEPES, glucose 5, Na pyruvate 5, MgCl_2_ 1.8, and CaCl_2_ 1, pH 7.4. The loaded cells were field-stimulated (5 V, 4 ms) at a frequency of 1 Hz. Sarcomere length (SL) and Fura-2 ratio (measured at 512 nm upon excitation at 340 nm and 380 nm) were simultaneously recorded using an IonOptix System (IonOptix, Westwood, Massachusetts, USA). Cell contractility was assessed by the percentage of sarcomere shortening. It is the ratio of twitch amplitude (difference of end-diastolic and peak systolic SL) to end-diastolic SL. Ca^2+^ transients were assessed by the percentage of variation of the Fura-2 ratio by dividing the twitch amplitude to end-diastolic ratio. The Tau parameter was used as an index of relaxation and Ca^2+^ transient decay kinetics. All parameters were calculated offline using a dedicated software (IonWizard 6x, Westwood, Massachusetts, Unites-States).

### Western Blot assay

Protein extracts were prepared from frozen cardiac tissue which were homogenized in a RIPA buffer containing: Tris HCl (pH 8.0) 50 mmol/L, NaCl 150 mmol/L, EDTA 2 mmol/L, NP40 1%, sodium deoxycholate acid 0.5%, SDS 0.1% supplemented with Complete Protease Inhibitor Tablets and PhosSTOPTM phosphatase inhibitor tablets (Roche Diagnostics, Basel, Switzerland). Tissue lysates were centrifuged at 15,000 g and 4°C for 20 min, pellets were discarded and supernatants were used. Protein samples were separated in denaturating acrylamide gels and subsequently transferred onto PVDF membranes. After blocking the membranes with 5% milk buffer for 1 h, the incubation with primary antibodies was carried out over night at 4°C. After incubation with the appropriate secondary antibody, proteins were visualized by enhanced chemiluminescence and quantified with ImageJ software. The primary antibodies used were: PDE2A blot on Figure 2 and supplementary Figure 1: goat anti-PDE2A, Santa Cruz, SC-17228; PDE2A blot on Figure 1 and 3: rabbit anti-PDE2A, Fabgennix, PDE2A-101AP; Vinculin: mouse anti-vinculin, V9131 Sigma-Aldrich; GAPDH: rabbit anti-GAPDH, cell signaling, 2118; FLAG: rabbit anti-flag L5, Novus biologicals, NBP1-06712.

**Figure 1.**
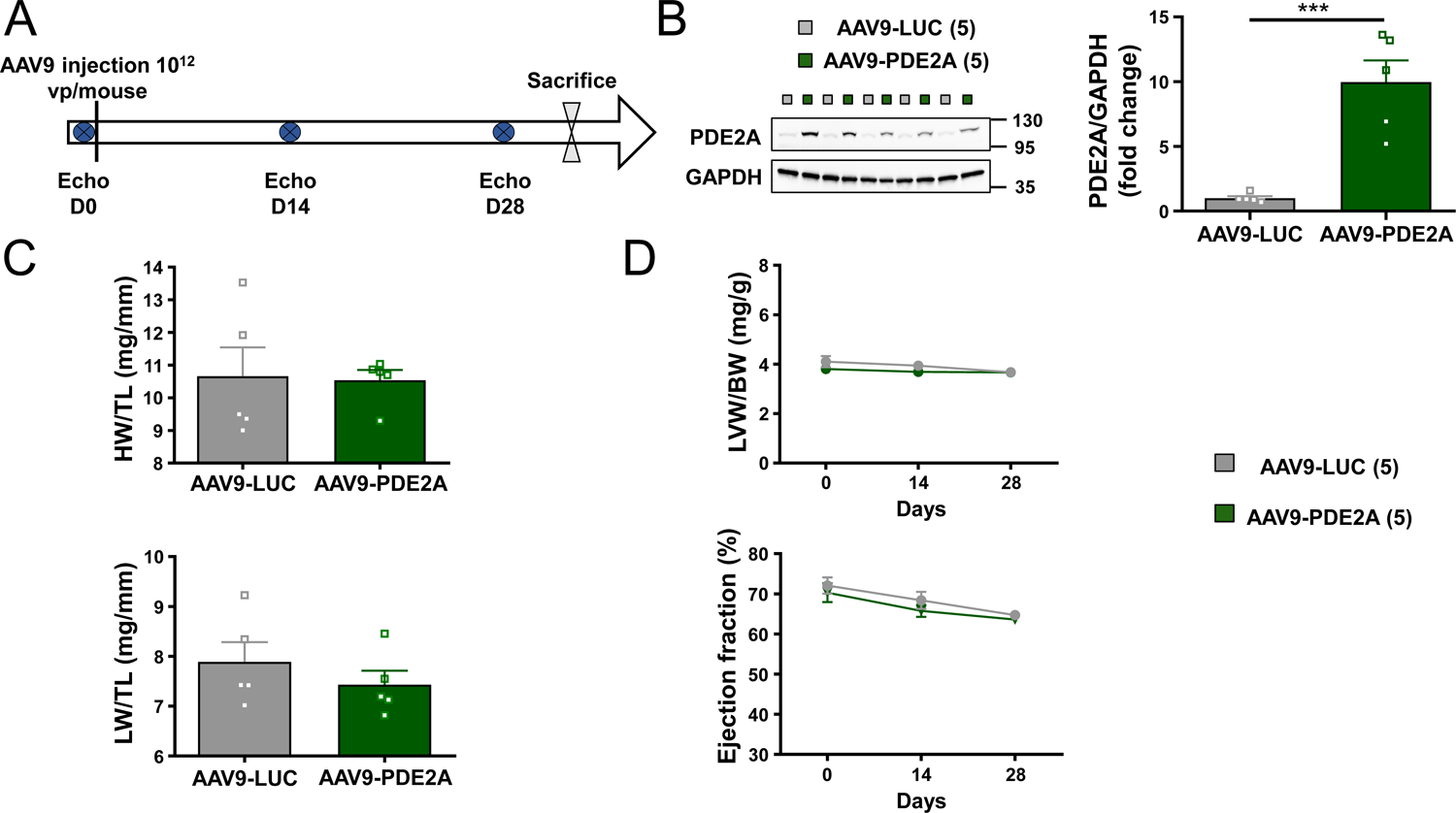
PDE2A overexpression by AAV9 does not alter cardiac function or morphological parameters. **A**, Schematic representation of the experimental protocol. Mice were injected in the tail vein with 10^12^ viral particles of a serotype 9 adeno-associated viruses encoding for Luciferase (AAV9-LUC) or phosphodiesterase 2A3 (AAV9-PDE2A). Twenty-eight days later, mice were sacrificed. Cardiac function was assessed throughout the protocol by echocardiography prior, then 2 and 4 weeks after the injection of the recombinant viruses. **B**, Left panel shows representative blots of PDE2A and GAPDH, the histogram represents the ratios of PDE2A over GAPDH quantified, expressed as mean ± SEM of fold change in cardiac tissues from AAV9-LUC and AAV9-PDE2A mice. **C**, Heart weight (HW) and lung weight (LV) over tibia length (TL) ratios expressed as mean ± SEM. **D**, Time course of ejection fraction and the ratio of calculated left ventricular weight (LVW) over body weight (BW) in AAV9-LUC and AAV9-PDE2A mice. Number of mice is indicated in the brackets. Statistical significance is indicated by *** p<0.001; Mann-Whitney (HW/TL) and Student’s t-tests (LW/TL) (B), two-way Anova followed by a Sidak test (D).

**Figure 2.**
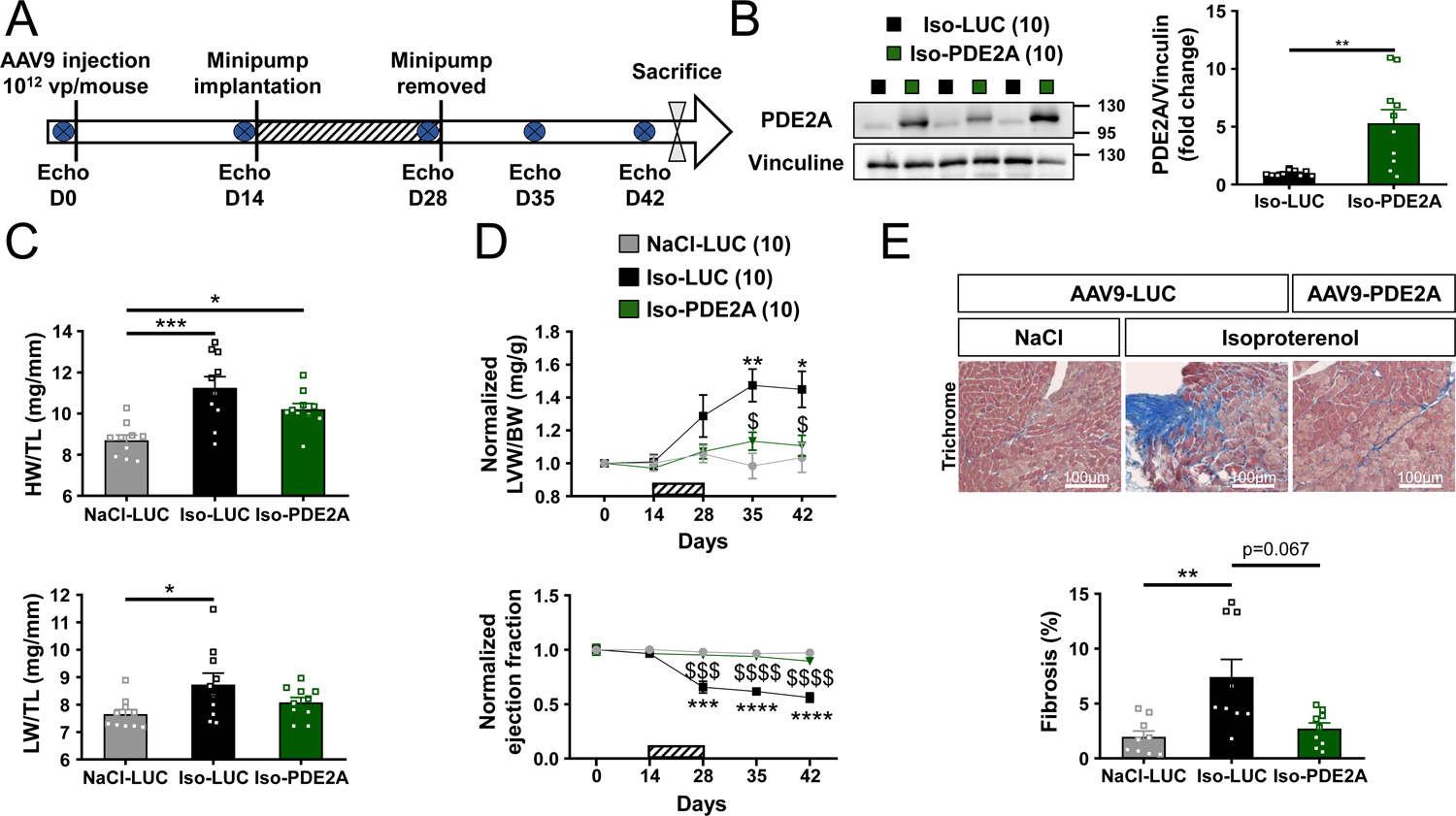
Gene therapy with PDE2A prevents cardiac remodeling and dysfunction induced by chronic isoproterenol infusion. **A**, Schematic representation of the experimental protocol. Mice were injected in the tail vein with 10^12^ viral particles of a serotype 9 adeno-associated viruses encoding for Luciferase (AAV9-LUC) or phosphodiesterase 2A3 (AAV9-PDE2A). Fourteen days later, mice were implanted subcutaneously with osmotic pumps diffusing either NaCl (NaCl-LUC) or isoproterenol at 60 mg/Kg/day (Iso-LUC or Iso-PDE2A). Treatment duration was 14 days (hatched bars). Minipumps were then removed and mice were kept for 2 additional weeks before sacrifice. Cardiac function was evaluated by serial echocardiography before injection of the virus as well as at 14, 28, 35, 42 days. **B**, Left panel shows representative blots of PDE2A and GAPDH. PDE2A protein expression in heart tissues extracts measured by Western blot and the histogram represents the ratios of PDE2A over GAPDH quantified expressed as mean ± SEM in NaCl-LUC, Iso-LUC and Iso-PDE2A groups. **C**, Heart weight (HW) and lung weight (LW) over tibia length (TL) ratio expressed as mean ± SEM. **D**, Time course of the normalized ratio of calculated left ventricular weight (LVW) over body weight (BW) and normalized ejection fraction in NaCl-LUC, Iso-LUC and Iso-PDE2A mice. Number of mice is indicated in the brackets. **E**, Panel on the left, representative images of Masson’s trichrome staining (scale bar 100 µm), histogram on the left represents quantifications of interstitial fibrosis as mean ± SEM in NaCl-LUC (n=9), Iso+Phe-LUC (n=9) and Iso+Phe-PDE2A (n=9) groups. * or $ p<0.05, ** or $$ p<0.01, *** or $$$ p<0.001, **** or $$$$ p<0.0001, * *vs*. NaCl-LUC and $ *vs*. Iso+Phe-LUC, Student’s t-test (B), Kruskal-Wallis followed by a Dunn’s test (C panel below and E), two-way Anova followed by a Tukey’s test (D), One-way Anova (C panel above) followed by a Tukey’s test.

**Figure 3.**
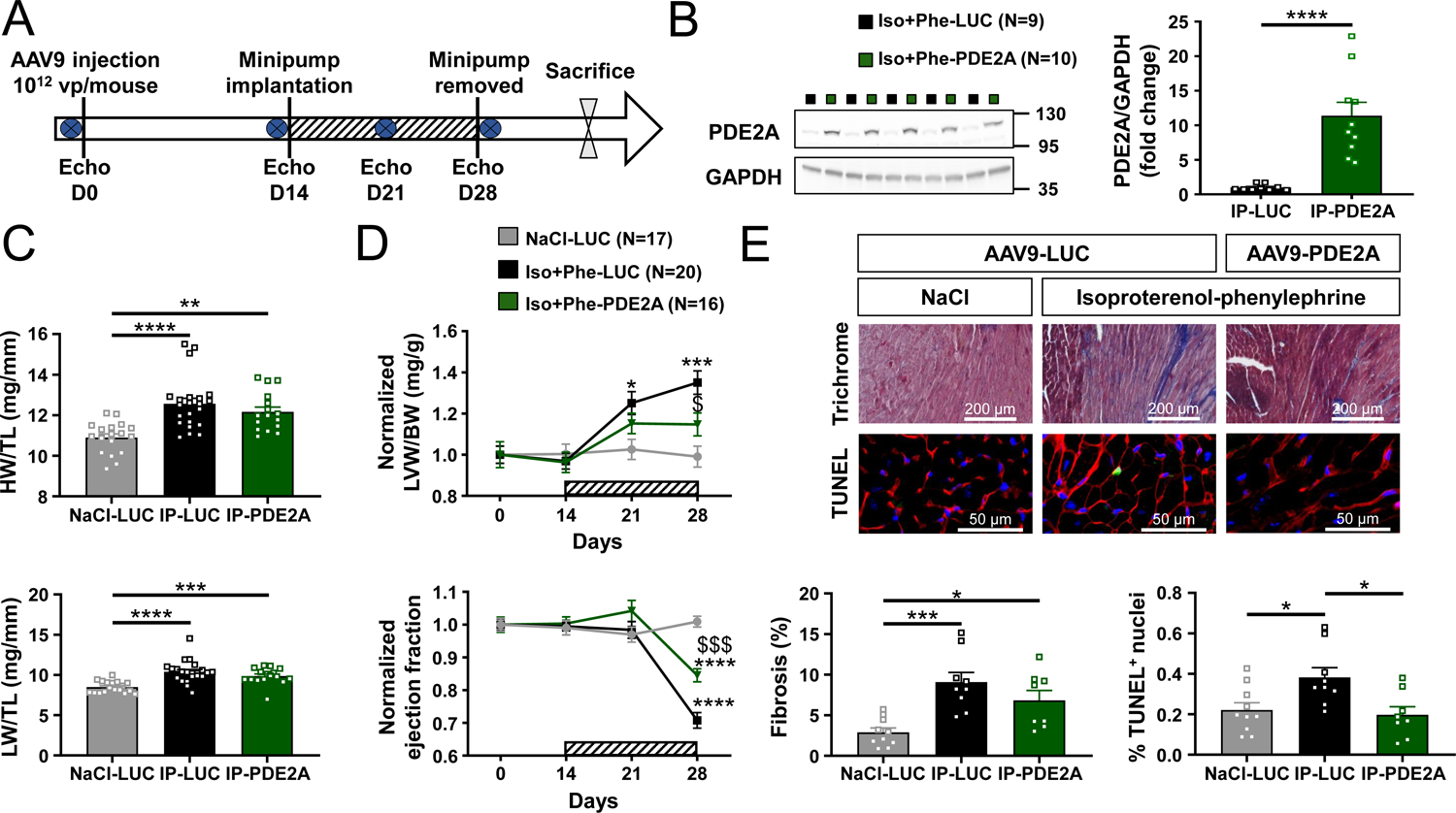
PDE2A3 overexpression limits maladaptive remodeling and cardiac dysfunction induced by chronic isoproterenol and phenylephrine infusion. **A**, Schematic representation of the experimental protocol. Mice were injected with 10^12^ viral particles of serotype 9 adeno-associated viruses (AAV9) encoding for Luciferase (AAV9-LUC) or phosphodiesterase 2A3 (AAV9-PDE2A). Fourteen days later, mice were implanted with osmotic minipumps diffusing either NaCl (NaCl-LUC) or isoproterenol+phenylephrine at 30 mg/Kg/day each (Iso+Phe-LUC and Iso+Phe-PDE2A). Minipumps were removed two weeks later and mice were sacrificed. Cardiac function was assessed throughout the protocol using echocardiography before the injection of the virus and at 2, 3, 4, 6 weeks. **B**, Representative blots of PDE2A and GAPDH on the left panel are shown and the histogram represents ratios of PDE2A over GAPDH quantifications expressed as Mean ± SEM in NaCl-LUC, Iso+Phe-LUC and Iso+Phe-PDE2A groups. **C**, Heart weight (HW) and lung weight (LW) over tibia length (TL) ratios expressed as mean ± SEM. **D**, Time course of the normalized ratios of calculated left ventricular weight (LVW) over body weight (BW) and normalized ejection fraction in NaCl-LUC, Iso+Phe-LUC and Iso+Phe-PDE2A mice. Number of mice is indicated in the brackets. **E**, Left panel shows representative images of Masson’s trichrome staining (scale bar 50 or 200 µm), the right panel depicts representative images of terminal deoxynucleotidyl transferase dUTP nick end labeling (TUNEL) assay (scale bar 100 µm) detecting apoptotic nuclei in green co-stained with the glycocalix marker wheat germ agglutinin in red and nuclei in blue colored with Hoechst. Bar graphs below representing mean ± SEM quantifications of interstitial fibrosis and apoptotic nuclei in NaCl-LUC (10), Iso+Phe-LUC (9) and Iso+Phe-PDE2A3 (9) mice are depicted. Statistical significance indicated by * or $ p<0.05, ** or $$ p<0.01, *** or $$$ p<0.001, **** or $$$$ p<0.0001, * *vs*. NaCl-LUC and $ *vs.* Iso+Phe-LUC was determined using Wilcoxon-Mann-Whitney test (B), Kruskal-Wallis followed by a Dunn’s test (C), Two-way Anova followed by a Tukey’s test (D), One-way Anova followed by a Tukey’s test (E).

### Quantitative real time polymerase chain reaction assay

Using Trizol reagent (MRCgene, Cincinnati, Ohio, USA), total RNA was extracted from ventricular tissue. Reverse transcription of RNA samples was carried out using iScript cDNA synthesis kit (Bio-Rad, Hercules, California, USA) according to manufacturer’s instructions. Real-time PCR reactions were prepared using SYBR Green Supermix (Bio-Rad, Hercules, California, USA) and performed in a CFX96 TouchTM Real-Time PCR Detection System (Bio-Rad, Hercules, California, USA). The relative amount of mRNA transcripts was quantified using the ΔC_t_ method. *Rplp2* and *Rpl32* housekeeping genes were used as reference for normalization. Sequences of the forward (For) and reverse (Rev) primers used for each studied gene are given below in the 5’-3’ orientation: *Pde2a* (For) TGGGGAACTCTTTGACTTGG, (Rev) ATGACCTTGCAGGAAAGCTG *Rplp2* (For) GCT GTG GCT GTT TCT GCT TC, (Rev) ATG TCG TCA TCC GAC TCC TC*Rpl32* (For) GCT GCT GAT GTG CAA CAA A, (Rev) GGG ATT GGT GAC TCT GAT GG

### Statistics

All data are presented as mean ± SEM. Normal distribution was tested by the Shapiro-Wilk normality test. An unpaired Student *t* test or paired Wilcoxon’s test were used for two group comparison. When data did not follow a normal distribution, a Mann-Whitney *U* test was used. Similarly, a one-way ANOVA was used to compare differences between multiple groups followed by a Tukey post hoc test, if a normal distribution of values was satisfied. If not, a Kruskal-Wallis test with Dunn’s post hoc test was employed. Artool two-way Mixed Anova Nested test or Barnard’s exact test were used when appropriate. A p value<0.05 was considered statistically significant.

## RESULTS

### Cardiac PDE2A3 overexpression in healthy C57BL/6 male mice does not alter the size, shape and function of the heart

Constructs including a CMV promotor to allow satisfactory expression of the PDE2A3 or of the luciferase (LUC) as an internal control of overexpression were built (supplemental Figure 1A). The AAV9-PDE2A3 promotes a cAMP-PDE activity which can be repressed by Bay 60-7550 (100 nmol/L), a specific PDE2 inhibitor, demonstrating that the adeno-associated virus allows to overexpress PDE2A specifically (supplemental Figure 1B). Prior to testing the therapeutic potential of enhanced cardiac PDE2A activity using engineered AAV9, we evaluated the effect of PDE2A3 overexpression on cardiac function and morphology in healthy mice. 10^12^ viral particles of AAV9-PDE2A or AAV9-LUC were injected to 8-week-old mice and followed up for 28 days (Figure 1A). As expected, cardiac PDE2A expression was significantly increased in AAV9-PDE2A3 mice with a ≈10-fold rise of PDE2A protein levels compared to AAV9-LUC mice measured by western blotting (Figure 1B). This overexpression neither affected the heart weight over tibia length ratio (HW/TL) nor the lung weight over tibia length ratio (LW/TL) (Figure 1C). Thus, increasing PDE2A activity in the heart did not provoke cardiac hypertrophy or lung congestion. LV morphology and function evaluated by transthoracic echocardiography remained unchanged with comparable LV weight over body weight ratio (LVW/BW) and ejection fraction (EF) measured in animals receiving AAV9-LUC and AAV9-PDE2A injections (Figure 1D and supplementary Table 1). Furthermore, heart rate measured in anesthetized mice as well as other morphological and echocardiographic parameters were not modified by PDE2A3 overexpression (supplementary Table 1).

### Cardiac gene therapy with PDE2A3 limits left ventricle maladaptive remodeling and dysfunction induced by a chronic infusion of catecholamines

Because the neurohormonal alterations of HF are characterized by marked elevations in norepinephrine and epinephrine,^29^ we hypothesized that PDE2A3 overexpression could exerts cardioprotective effects in two different mouse models of cardiac dysfunction evoked by chronic infusion with catecholamines. In both trials, tail vein injections with 10^12^ viral particles of AAV9-LUC or AAV9-PDE2A were performed 14 days prior to minipump implantation. We first surmised that increased PDE2A activity may mitigate left ventricular (LV) remodeling and dysfunction in response to a chronic β-AR stimulation. Thus, we chose to subject the mice to a 14 day-infusion with the non-selective β-AR agonist Iso (60 mg/Kg/day),^17, 30^ then the osmotic pumps were removed. Cardiac function was evaluated *in vivo* by transthoracic echocardiography over 6 weeks (Figure 2A). Analysis of PDE2A mRNA levels (supplementary Figure 2) and protein expression revealed a ≈6-fold increase in the harvested hearts from mice injected with AAV9-PDE2A3 (Figure 2B). Both groups of mice expressing either LUC or PDE2A3 implanted with Iso minipumps displayed a similar increase in HR at 2 weeks, attesting the adequate infusion of Iso (supplementary Figure 3A). This treatment resulted in a marked increase of HW/TL ratio indicating cardiac hypertrophy (P<0.001), accompanied with lung congestion (p<0.05) (Figure 2C). Echocardiography confirmed LV hypertrophy and revealed a significantly decreased EF after 2 weeks’ infusion with Iso (Figure 2D). This LV hypertrophy was accompanied with increased end systolic diameter (LVIDs) evoking contractile dysfunction, and augmented diastolic LV diameter (LVIDd) suggesting left ventricular dilation (supplementary Table 2). Moreover, Masson’s Trichrome staining revealed a significant increase in fibrosis after the chronic Iso treatment (p<0.01; Figure 2E). While the HW/TL ratio and congestion were only slightly ameliorated in mice overexpressing PDE2A (Figure 2C), LV hypertrophy, systolic dysfunction, myocardial fibrosis and LV dilation induced by Iso were totally prevented (Figure 2 and supplementary Table 2).

Since elevated catecholamines in HF activate not only β-AR but also α-adrenoceptors (α-AR), we added phenylephrine (Phe) to Iso in the minipumps at 30 mg/kg/day each, an infusion which better recapitulates the alterations observed in pressure overload-induced experimental HF and in human hypertrophic cardiomyopathy^31^ (Figure 3A). In this set of experiments, a ≈12-fold increase of PDE2A3 expression in the ventricular tissue was achieved with AAV9 (Figure 3B). Chronic infusion with Iso+Phe induced cardiac hypertrophy and lung congestion as attested by a significant increase in HW/TL and LW/TL ratios (p<0.001; Figure 3C). These effects were not prevented by PDE2A3 overexpression despite a trend to diminution (Figure 3C). However, echocardiography revealed that LV hypertrophy and dysfunction induced by β-AR and α-AR stimulation were significantly attenuated in AAV9-PDE2A3 mice (p<0.05 and p<0.001 *vs.* Iso+Phe-LUC; Figure 3D), despite a persistent left ventricular chamber dilation in diastole and systole accompanied with an increased posterior wall thickness in diastole (p<0.05; supplementary Table 3). While fibrosis was not significantly reduced by PDE2A3 overexpression, apoptosis assessed by the TUNEL assay was clearly diminished (p<0.05; Figure 3E). Also, gene therapy with PDE2A3 preserved the diastolic function of the treated mice. Indeed, mice treated with Iso+Phe had a clear diminished transmitral blood flow during the left atria (LA) contraction phase as indicated by the significant decrease of the A wave (p<0.0001; supplementary Table 3) analyzed with pulse wave Doppler. This suggests that chronic Iso+Phe treatment altered LA contractility or compliance, explaining the trend to increased E/A ratio noticed (p=0.07, supplementary Table 3). These alterations of the diastolic function induced by catecholamines were prevented by the gene therapy with PDE2A3 (supplementary Table 3).

### PDE2A3 improves defective β-AR responses of isolated cardiomyocytes after chronic catecholamines infusion

To investigate how gene therapy with PDE2A3 affects the ECC and its β-AR modulation at the cellular level, we measured sarcomere shortening (SS) and calcium transients (CaT) simultaneously in Fura-2–loaded ventricular myocytes (VMs) paced at 1 Hz. As shown in Figure 4A-D, baseline CaT and SS amplitudes and time constants for relaxation (τ) were not modified in VMs from Iso+Phe treated mice although CaT decay was slower in cells overexpressing PDE2A3 (p<0.01 *vs.* Iso+Phe-LUC, Figure 4B). When challenged with a submaximal concentration of Iso (3 nmol/L), VMs isolated from NaCl-LUC mice had significantly increased CaT and SS amplitudes by 1.9-3.5-fold respectively (p<0.0001). Iso also strongly accelerated the relaxation rates of both CaT and SS (p<0.0001 *vs.* Ctrl, Figure 4D). These positive inotropic and lusitropic effects were significantly blunted in VMs isolated from LUC mice treated with Iso+Phe. Indeed, Iso had no significant effect on CaT amplitude and kinetics while SS amplitude only increased by 1.6-fold (p=0.0002, *vs*. NaCl-LUC at baseline; Figure 4A-D). Interestingly, the positive inotropic effects of Iso were partially restored in VMs from PDE2A3-overexpressing animals, although decay times were not improved. Thus, these results demonstrate that gene therapy with PDE2A3 improves β-AR responsiveness of VMs dampened by chronic infusion with catecholamines in mice.

**Figure 4.**
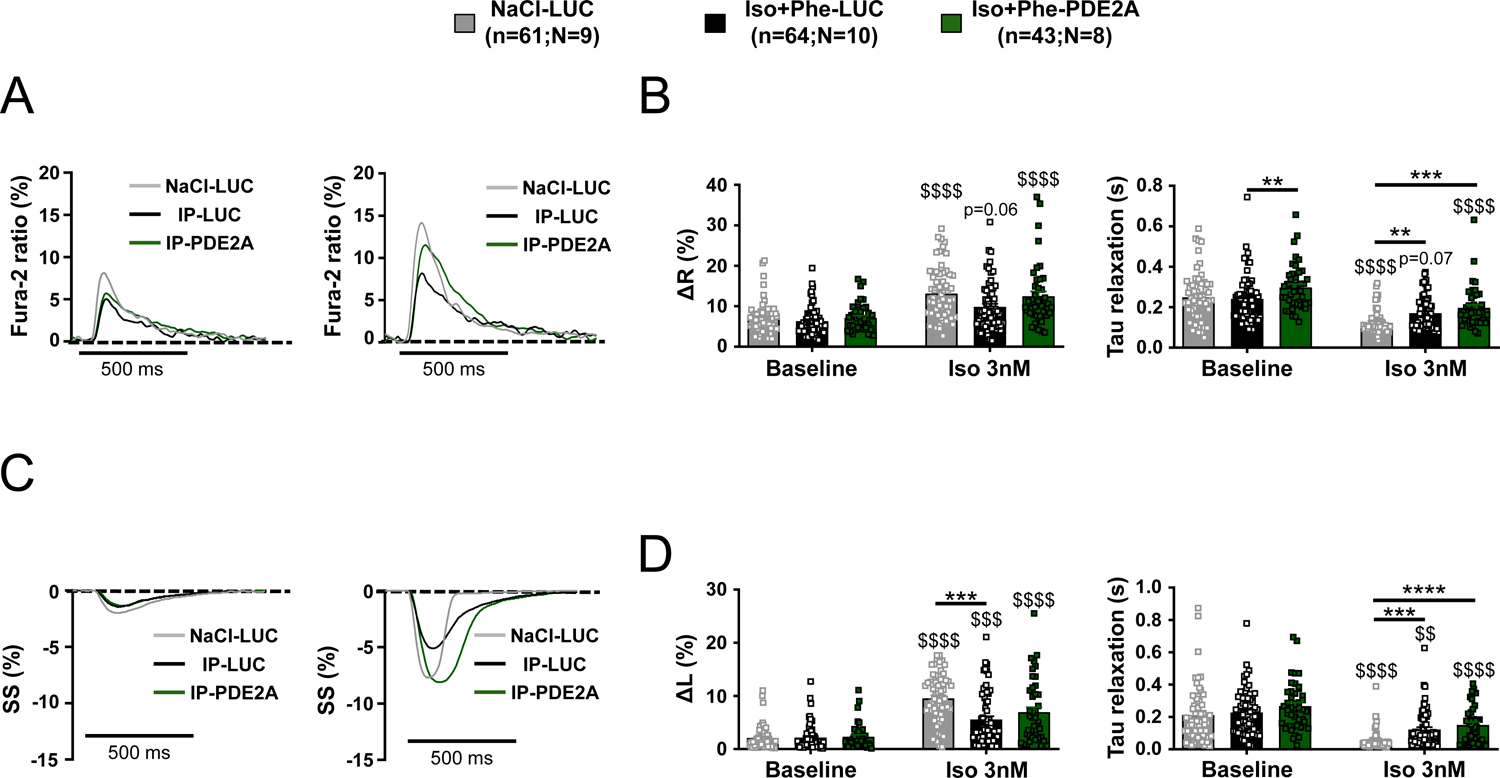
PDE2A overexpression ameliorates the β-AR responsiveness of the excitation-contraction coupling. **A**, Representative traces of calcium transients (CaT) and **C**, sarcomere shortening (SS) obtained simultaneously from ventricular myocytes loaded with Fura-2 and paced (1 Hz) isolated from NaCl-LUC (grey), Iso+Phe-LUC (black) or Iso+Phe-PDE2A (green) mice in control conditions and in the presence of 3 nmol/L isoproterenol (Iso). **B**, Representative histograms of mean ± SEM of the CaT amplitudes (expressed as the percentage of diastolic Fura-2 ratio) (Left panel) and of the kinetics for return to Ca^2+^ diastolic levels (right panel) in control conditions and in the presence of 3 nmol/L isoproterenol. **D**, Representative histograms of mean ± SEM of SS amplitude expressed as the percentage of resting sarcomere length (left panel) and of relaxation time constant (tau, right panel) at baseline and in the presence of 3 nmol/L Iso. **D**, SS and CaT were measured in 43 to 64 cells from 9 NaCl-LUC (grey bars), 10 Iso+Phe-LUC (black bars) and 8 Iso+Phe-PDE2A (green bars) mice. Numbers are indicated in the brackets as following (n=number of cells; N=Number of mice). **** or $$$$ p<0.0001, *** or $$$ p<0.001, ** or $$ p<0.01, * or $ p<0.05 ($ *vs.* corresponding group at baseline); Artool two-way mixed Anova nested test (B, D).

### PDE2A3 cardiac overexpression prevents spontaneous calcium events and ventricular arrhythmias

It has been previously reported that a constitutive cardiac overexpression of PDE2A3 had antiarrhythmic effects.^19, 32^ We thus tested whether similar effects could be produced by an acute overexpression.

First, we investigated the occurrence of pro-arrhythmogenic SCEs recorded in VMs isolated from the three groups of mice stimulated with a maximal Iso concentration of 100 nmol/L (Figure 5A). The time to first SCE and the percentage of cells presenting SCEs in each group were similar (supplementary Figure 4B). However, the number of SCEs recorded during a 3 min period of maximal β-AR stimulation was increased by Iso 100 nmol/L in the Iso+Phe-LUC group, but these cellular pro-arrhythmogenic events were nearly absent in cells isolated from Iso+Phe-PDE2A group (Figure 5B).

**Figure 5.**
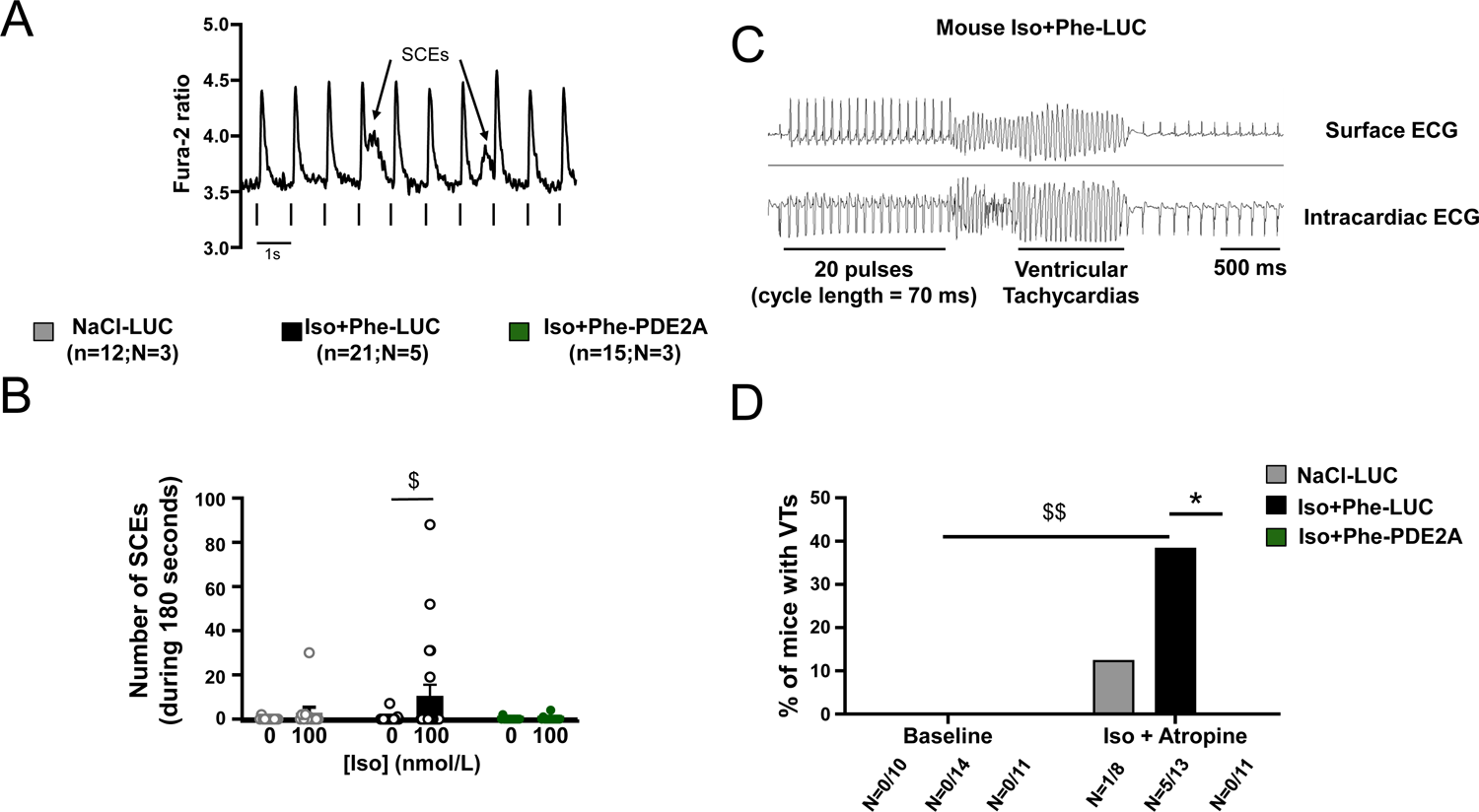
Gene therapy with PDE2A3 exerts anti-arrhythmic effects at the cellular level and *in vivo*. **A**, Representative traces of spontaneous calcium release events pointed by arrow, expressed as the percentage of diastolic Fura-2 ratio in paced (1Hz) ventricular myocytes isolated from Iso+Phe-LUC mice in the presence of 100 nmol/L isoproterenol. **B**, SCEs were counted during the 3 min following isoproterenol 100 nmol/L administration. SCEs were measured in 12 to 21 cells isolated from 3 NaCl-LUC (grey), 5 Iso+Phe-LUC (black) and 3 Iso+Phe-PDE2A (green) mice. Numbers are indicated in the brackets as following (n=number of cells; N=number of mice). **C**, Representative examples of simultaneous lead 1 ECG and intraventricular electrogram recordings obtained in a mouse subjected to gene therapy with AAV9-LUC and chronically infused with isoproterenol and phenylephrine (Iso+Phe, 30 mg/kg/day each) for 14 days after paced extrasystoles. The protocol consisted of twenty pulses at a cycle length of 70 ms followed by one then two then three closer pulses (cycle length determined by the refractory period). Ventricular tachycardias (VTs) were defined as the occurrence after the last paced beat of at least 4 consecutive QRS complexes with a different morphology from that seen in normal sinus rhythm. **D**, Histogram showing the percentage of mice with arrhythmias in all three groups (NaCl-LUC, Iso+Phe-LUC, Iso+Phe-PDE2A3) at baseline and after isoproterenol 1.5 mg/kg + atropine 1 mg/kg injection. The number of animals presenting VTs per group is indicated beneath the histogram. * or $ p<0.05 ($ *vs.* corresponding group at baseline); Wilcoxon test’s test ($ *vs.* baseline) (B); Barnard’s exact test * or $ p<0.05 ($ *vs.* corresponding group at baseline) (D).

Next, we tested the putative antiarrhythmic effect of PDE2A gene therapy *in vivo*. We evaluated the propensity of the treated animals to develop ventricular tachycardia. To do so, intracardiac recordings and pacing were performed. Six-lead ECGs recorded in baseline conditions revealed similar PR interval, QRS complex and QTc interval durations in NaCl-LUC, Iso+Phe-LUC and Iso+Phe-PDE2A3 animals (supplementary Table 4). However, Iso+Phe-LUC mice had a significantly lower RR interval (p<0.001 *vs.* NaCl), thus higher heart rate (HR, p<0.001 *vs.* NaCl). Interestingly, despite their similar treatment with catecholamines, mice injected beforehand with AAV9-PDE2A3 had a cardiac frequency equivalent to that of control animals (supplementary Table 4). Mice were then subjected to catheter-mediated ventricular pacing in baseline conditions which did not evoke cardiac arrhythmias in any animal of the 3 groups. Intraperitoneal injections of Iso+atropine (1.5 mg/kg and 1 mg/kg, respectively) were therefore performed to favor ventricular tachycardia (VT) occurrence. As expected, in the three groups of animals, injection of Iso+atropine significantly increased HR, decreased RR and PR intervals, increased QRS complex durations (but not significantly in NaCl-LUC mice), while QTc duration remained stable (supplementary Table 4). While LUC mice treated or not with catecholamines and animals overexpressing PDE2A3 had similar ECG parameters following the Iso+atropine injection, they responded differently to ventricular pacing. Indeed, extra-stimuli delivered following trains of 20 impulses triggered arrhythmias in only 1 over 8 NaCl-LUC mice but VT episodes as depicted in Figure 5C were evoked in 5 out of 13 LUC animals chronically infused with catecholamines (p<0.05 *vs* baseline). However, none of the 11 mice overexpressing PDE2A3 had arrhythmias when applying extrasystolic stimuli in a programmed electrical stimulation (PES) protocol (Figure 5D). This demonstrates that gene therapy with PDE2A3 protects from ventricular arrhythmias upon stress conditions. To test whether these antiarrhythmic effects are specific to PDE2A overexpression, we evaluated the potency of PDE4B3 to achieve such protective effects, since we previously reported that gene therapy with this enzyme prevents adverse remodeling in HF mice models.^17^ Interestingly, a similar treatment with an AAV9 encoding for PDE4B3 did not produce similar anti-arrhythmic effects despite diminished LV hypertrophy and dysfunction (supplementary Figure 5A-C). This demonstrates that protection against VTs evoked by catecholamines is better achieved with an increase of PDE2A activity than PDE4B3 in this model.

## DISCUSSION

Clinically, PDE inhibition has been considered a promising approach to compensate for the catecholamine desensitization that accompanies HF. In that respect, PDE3 inhibitors, such as milrinone or enoximone, have been used clinically to improve systolic function and alleviate the symptoms of acute HF. However, their chronic use has proven to be detrimental, increasing adverse remodeling^33^ and ventricular arrhythmias.^34, 35^ These adverse effects could potentially be avoided using compartment-specific, PDE isozyme-selective inhibitors.^10^ Here, we proposed to test the opposite strategy, *i.e.* increasing rather than inhibiting PDE activity. We believe that this strategy, which is reminiscent of the counter-intuitive beneficial effect of beta-blockers in HF, could be therapeutically relevant in HF because it would prevent a deleterious accumulation of cAMP during catecholamines spill over. In that line, we recently demonstrated that constitutive overexpression of PDE4B,^17^ one of the main PDE4 isoform expressed in the cardiomyocyte to control the β-AR regulation of the ECC,^36^ is cardioprotective. We also showed that gene therapy with AAV9-PDE4B exerts cardioprotective effects limiting adverse remodeling evoked by catecholamines or increased postcharge.^17^ Similarly, PDE2A constitutive overexpression exerts anti-hypertrophic effects^18^ and transgenic mice overexpressing PDE2A have preserved ejection fraction after myocardial infarction and are protected against catecholamine induced ventricular arrhythmia.^19^ It was thus important to test whether an acute increase in PDE2A activity, as obtained during AAV9 gene transfer, could be therapeutically relevant.

Similarly to what we reported earlier in the transgenic mice,^19^ here we found that acute overexpression of PDE2A obtained after systemic injection of the AAV9 construct did not produce any noticeable change neither on the left ventricle size nor its function. Morphometric parameters were not impacted by the overexpression of PDE2A, and contractile parameters were preserved with no congestion.

The safety of this treatment allowed us to move forward and test whether overexpression of PDE2A would counteract the adverse remodeling evoked by catecholamines. Indeed, AAV9 delivery of PDE2A efficiently protected against the detrimental effects of chronic isoproterenol treatment. Chronic catecholamine infusion models promote cardiomyocyte death and contractile cells are then replaced by interstitial fibrosis.^37^ AAV9-mediated PDE2A overexpression counteracted the systolic dysfunction, attenuated the hypertrophic response, and efficiently prevented fibrosis, a hallmark of pathological remodeling. These cardioprotective effects against β-AR chronic stimulation were as effective as the one obtained by gene therapy with PDE4B,^17^ confirming that increasing PDE activity is effective to counteract adverse remodeling evoked by catecholamines. However, elevated catecholamines act not only via β-AR but also via α-AR. Interestingly, gene therapy with PDE2A (and PDE4B, supplemental Figure 5) was able to limit LV hypertrophy and dysfunction induced by Iso+Phe challenge, a situation which recapitulates more effectively early transcriptional alterations observed in pressure overload-induced experimental HF and in human hypertrophic cardiomyopathy.^31^ Cardioprotective effects of PDE2A overexpression were less pronounced with Iso+Phe than with Iso alone, probably because PDE2A is less efficient in counteracting cAMP-independent effects of α_1_-AR stimulation such as apoptosis and fibrosis,^31^ than cAMP-signalling emanating from β-AR stimulation. Nevertheless, adverse remodeling and LV dysfunction were reduced by PDE2A overexpression.

Our results demonstrate that gene therapy with PDE2A improves cardiac adaptation to catecholaminergic stress. Indeed, we found that PDE2A cardiac overexpression led to a better responsiveness of the ECC to the β-AR agonist which could be explained by decreased intracellular cAMP levels upon chronic elevation of catecholamines known to promote heterologous desensitization of β-AR by promoting their phosphorylation by PKA.^38^ This may contribute to the improved ejection fraction we observed in animals subjected to gene therapy with PDE2A. Furthermore, we observed less pro-arrhythmogenic spontaneous calcium events in cells isolated from PDE2A overexpressing hearts, similarly to what has been described recently in cells from PDE2A transgenic mice.^32^ This phenomenon could also contribute to improve the cardiac function and decrease the propensity of these animals to trigger ventricular arrhythmias as we unveil here, in accordance with the protection conferred by PDE2A constitutive overexpression in the setting of myocardial infarction previously described.^19^

While we showed that PDE2A overexpression is cardioprotective,^18^ another group showed that inhibition rather than increased activity of PDE2 exerts positive outcome in a pressure-overload model.^26^ Furthermore, under pathologic conditions PDE2A overexpression was proposed to increase ventricular cardiomyocyte size, thus to evoke adverse remodeling.^24^ Although it was demonstrated that PDE2 regulates cGMP generated by particulate guanylate cyclases,^39^ the cardioprotective effects of PDE2 inhibition were attributed to increased cGMP emanating from the NO stimulated soluble guanylate cyclase.^26^ However, PDE2 inhibition by promoting voltage gated Ca^2+^ transients in postganglionic neurons from stellate ganglia increases cardiac neurotransmission,^40^ a mechanism which may also contribute to increase cardiac function and alleviate cardiac dysfunction. Moreover, global PDE2 inhibition induced a modest coronary vasodilatation,^26^ which could have contributed to the beneficial effects observed.

Here, using AAV9, we target preferentially the heart, and we believe that cardioprotective effects are achieved due to this preferential cardiac overexpression of PDE2. Interestingly, activating mutations in another PDE, PDE3, responsible for a rare disease characterized by the combination of brachydactyly and hypertension, confers cardioprotection despite the increase in afterload.^41^ This latter observation supports our initial postulate and our observations reported here and earlier,^17^ that increasing rather than decreasing the activity of PDEs in cardiac tissue is beneficial. However, it remains to be demonstrated which PDE will be the most effective to achieve this goal. We show here that gene therapy with PDE2A is as effective as PDE4B in counteracting the adverse remodelling of the left ventricle and systolic dysfunction induced by chronic catecholamines infusion, but less efficient in counteracting stress-induced ventricular tachycardia. This difference may come from the intrinsic characteristics of the two enzymes. Indeed, PDE2A has a 20-fold lower affinity for cAMP than PDE4B (K_m_=30 *vs.* 1.5–4.7 µmol/L),^42^ and a ∼1000-fold higher hydrolytic activity (V_max_=120 *vs.* 0.13 µmol/min/mg protein).^42^ Thus, PDE2A would only be turned on when cAMP concentration reaches supra-physiological values, and its high V_max_ would tend to rapidly reduce it to normal. In other words, PDE2A might serve as a *safety valve* to avoid the cell to be flooded with cAMP. In contrast, PDE4B degrades cAMP at physiological concentrations and, as such, plays a more essential role in the regulation of ECC.^36^ Its overexpression will thus have different consequences on heart function, notably by reducing the β-AR response of the heart.

In conclusion, our results demonstrate that increasing PDE2A activity is beneficial in heart failure. They suggest that a gene therapy with PDE2A in cardiac cells, or PDE2A activators yet to be discovered, could be of therapeutic value. PDE2A is a cGMP-activated PDE^43^ and its cAMP-hydrolytic activity can be increased ∼30 fold upon cGMP binding to its GAFB domain. cGMP elevation, which has been shown to be cardioprotective in HF,^44^ could be achieved by activating the particulate guanylate cyclase localized near PDE2A,^45^ to constitute a NP/BNP/cGMP-triggered defence mechanism during cardiac stress, in particular during excessive β-AR drive. Interestingly, sacubitril which further elevates NPs improves classical treatments for HF,^46^ and it has been recently demonstrated that NPs exert antiarrhythmic effects via PDE2.^23^ We can therefore speculate that further beneficial effects could be obtained with a gene therapy with PDE2 which could constitute a complement to neprilysin inhibitors and actual treatments of chronic HF to limit adverse remodelling and accompanying arrhythmias.

